# Afferent-efferent connectivity between auditory brainstem and cortex accounts for poorer speech-in-noise comprehension in older adults

**DOI:** 10.1101/568840

**Authors:** Gavin M. Bidelman, Caitlin N. Price, Dawei Shen, Stephen R. Arnott, Claude Alain

## Abstract

Age-related hearing loss leads to poorer speech comprehension, particularly in noise. Speech-in-noise (SIN) deficits among the elderly could result from weaker neural activity within, or poorer signal transmission between brainstem and auditory cortices. By recording neuroelectric responses from brainstem (BS) and primary auditory cortex (PAC), we show that beyond simply attenuating neural activity, hearing loss in older adults compromises the transmission of speech information between subcortical and cortical hubs of the auditory system. The strength of afferent BS→PAC neural signaling (but not the reverse efferent flow; PAC→BS) varied with mild declines in hearing acuity and this “bottom-up” functional connectivity robustly predicted older adults’ SIN perception. Our neuroimaging findings underscore the importance of brain connectivity, particularly afferent neural communication, in understanding the biological basis of age-related hearing deficits in real-world listening environments.

## INTRODUCTION

Difficulty perceiving speech in noise (SIN) is a hallmark of aging. Hearing loss and reduced cognitive flexibility may contribute to speech comprehension deficits that emerge after the fourth decade of life (Humes, 1996; Humes et al., 2012). Yet, older adults’ SIN difficulties persist even without substantial hearing impairments (Gordon-Salant and Fitzgibbons, 1993; Schneider et al., 2002), suggesting robust speech processing requires more than audibility.

Emerging views of aging suggest that in addition to peripheral changes (i.e., cochlear pathology) (e.g., Chambers et al., 2016), older adults’ perceptual SIN deficits might arise due to poorer sensory encoding, *transmission*, and decoding of acoustic speech features within the brain’s central auditory pathways (Schneider et al., 2002; Wong et al., 2010; Peelle et al., 2011; Anderson, White-Schwoch, et al., 2013a). Although “central presbycusis” offers a powerful framework for studying the perceptual consequences of aging (Humes, 1996), few studies have explicitly investigated how the auditory system extracts and transmits features of the speech signal across different levels of the auditory neuroaxis. Senescent changes have been observed in pontine, midbrain, and cortical neurons (Peelle and Wingfield, 2016). Yet, such insight into brainstem-cortex interplay has been limited to animal models (Chambers et al., 2016).

Age-related changes in hierarchical auditory processing can be observed in scalp-recorded frequency-following responses (FFR) and event-related brain potentials (ERPs), dominantly reflecting activity of midbrain and cerebral structures, respectively (Bidelman et al., 2013). Both speech-FFRs (Anderson, White-Schwoch, et al., 2013b; Bidelman, Villafuerte, et al., 2014) and ERPs (Tremblay et al., 2003; Alain et al., 2014; Bidelman, Villafuerte, et al., 2014) reveal age-related changes in the responsiveness (amplitude) and precision (timing) of how subcortical and cortical stages of the auditory system extract complex sounds. In our studies recording these potentials simultaneously, we showed aging is associated with increased redundancy (higher shared information) between brainstem and cortical representations for speech (Bidelman, Villafuerte, et al., 2014; Bidelman et al., 2017). Our previous findings imply that SIN problems in older listeners might result from aberrant *transmission* of speech signals from brainstem en route to auditory cortex, a possibility that has never been formally tested.

A potential candidate for these central encoding/transmission deficits in aging (Humes, 1996) could be the well-known afferent and efferent (corticofugal) projections that carry neural signals bidirectionally between brainstem and primary auditory cortex (BS↔PAC) (Suga et al., 2000; Bajo et al., 2010). Descending corticocollicular (PAC→BS) fibers have been shown to calibrate sound processing of midbrain neurons by fine tuning their receptive fields in response to behaviorally relevant stimuli (Suga et al., 2000). Germane to our studies, corticofugal efferents drive learning-induced plasticity in animals (Bajo et al., 2010) and may also account for the neuroplastic enhancements observed in human FFRs across the age spectrum (Musacchia et al., 2007; Wong et al., 2007; Anderson, White-Schwoch, et al., 2013b). While assays of olivocochlear (peripheral efferent) function are well-established (e.g., otoacoustic emissions; de Boer and Thornton, 2008) there have been no direct measurements of corticofugal (central efferent) system function in humans, despite its assumed role in complex listening skills like SIN (Slee and David, 2015).

To elucidate brainstem-cortical reciprocity in humans, we recorded neuroelectric FFR and ERP responses during active speech perception. Examining older adults with normal or mild hearing loss for their age allowed us to investigate how hierarchical coding is changed with declining sensory input. We used source imaging and functional connectivity analyses to parse activity *within* and directed (causal) transmission *between* sub- and neo-cortical levels. To our knowledge, this is the first study to document afferent and corticofugal efferent function in human speech processing. We hypothesized (i) hearing loss would alter the relative strengths of afferent (BS→PAC) and/or corticofugal (PAC→BS) signaling and more importantly, (ii) poorer connectivity would account for older adults’ perceptual SIN deficits. Beyond aging, such findings would also establish a biological mechanism to account for the pervasive, parallel changes in brainstem and cortical speech-evoked responses previously observed in highly skilled listeners (e.g., musicians) and certain neuropathologies (Musacchia et al., 2008; Bidelman and Alain, 2015; Bidelman et al., 2017).

## METHODS

### Participants

Thirty-two older adults aged 52-75 years were recruited from the Greater Toronto Area to participate in our ongoing studies on aging and the auditory system. None reported history of neurological or psychiatric illness. Pure-tone audiometry was conducted at octave frequencies between 250 and 8000 Hz. Based on listeners’ hearing thresholds, the cohort was divided into normal and hearing-impaired groups (Fig. 1A). In this study, normal-hearing (NH; *n*=13) listeners were classified as having average thresholds (250 to 8000 Hz) better than 25 dB HL across both ears, whereas listeners with hearing loss (HL*; n*=19) had average thresholds poorer than 25 dB HL. This division resulted in pure-tone averages (PTAs) (i.e., mean of 500, 1000, 2000 Hz) that were ~10 dB better in NH compared to HL listeners (*mean ±SD*; NH: 15.3±3.27 dB HL, HL: 26.4±7.1 dB HL; *t*_2.71_=-5.95, *p*<0.0001). This definition of hearing impairment further helped the post hoc matching of NH and HL listeners on other demographic variables while maintaining adequate sample sizes per group. Both groups had signs of age-related presbycusis at very high frequencies (8000 Hz), which is typical in older adults. However, it should be noted that the audiometric thresholds of our NH listeners were better than the hearing typically expected based on the age range of our cohort, even at higher frequencies (Pearson et al., 1995; Cruickshanks et al., 1998).

**Figure 1:**
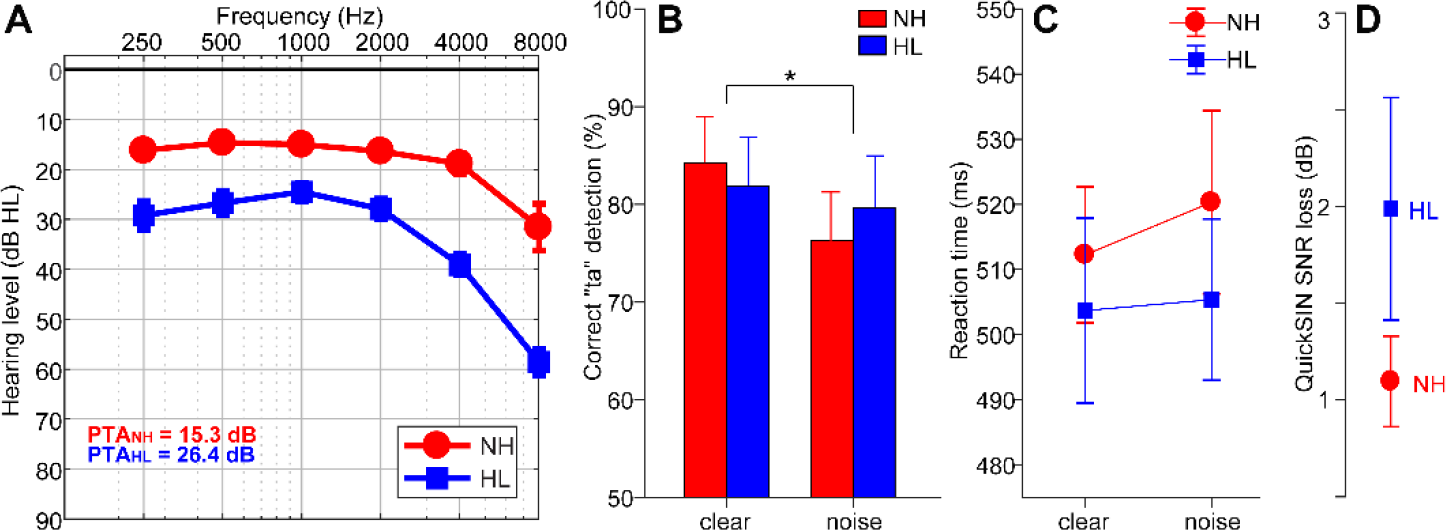
Audiometric and perceptual results. (A) Audiograms for listeners with normal hearing (NH) and hearing loss (HL). Hearing was ~10 dB better in NH vs. HL listeners. (B) Behavioral accuracy for detecting infrequent /ta/ tokens in clear and noise-degraded conditions. Noise-related declines in behavioral performance were prominent but no group differences were observed. (C) Reaction times (RTs) for speech detection were similar between groups and speech SNRs. (D) HL listeners showed more variability and marginally poorer QuickSIN performance than NH listeners. errorbars = ± s.e.m., \**p*< 0.05.

Importantly, besides hearing, the groups were otherwise matched in age (NH: 66.2±6.1 years, HL: 70.4±4.9 years; *t*_2.22_=-2.05, *p* = 0.052) and gender balance (NH: 5/8 male/female; HL: 11/8; Fisher’s exact test, *p*=0.47). Age and hearing loss were not correlated in our sample (Pearson’s *r*=0.29, *p*=0.10). Participants were compensated for their time and gave written informed consent in compliance with a protocol approved by the Baycrest Centre research ethics committee.

### Stimuli and task

Three tokens from the standardized UCLA version of the Nonsense Syllable Test were used in this study (Dubno and Schaefer, 1992). These tokens were naturally produced English consonant-vowel phonemes (/ba/, /pa/, and /ta/), spoken by a female talker. Each phoneme was 100-ms in duration and matched in terms of average root mean square sound pressure level (SPL). Each had a common voice fundamental frequency (F0=150 Hz) and first and second formants (F1= 885, F2=1389 Hz). This relatively high F0 ensured that FFRs would be of dominantly subcortical origin and cleanly separable from cortical activity (Bidelman, 2018), since PAC phase-locking (cf. “cortical FFRs”; Coffey et al., 2016) is rare above ~100 Hz (Brugge et al., 2009; Bidelman, 2018). CVs were presented in both clear (i.e., no noise) and noise-degraded conditions. For each noise condition, the stimulus set included a total of 3000 /ba/, 3000/pa/, and 210 /ta/ tokens (spread evenly over three blocks to allow for breaks).

For each block, speech tokens were presented back-to-back in random order with a jittered interstimulus interval (95-155 ms, 5ms steps, uniform distribution). Frequent (/ba/, /pa/) and infrequent (/ta/) tokens were presented according to a pseudo-random schedule such that at least two frequent stimuli intervened between target /ta/ tokens. Listeners were asked to respond each time they detected the target (/ta/) via a button press on the computer. Reaction time (RT) and detection accuracy (%) were logged.

These procedures were then repeated using an identical speech triplet mixed with eight talker noise babble (cf. Killion et al., 2004) at a signal-to-noise ratio (SNR) of 10 dB. Thus, in total, there were 6 blocks (3 clear, 3 noise). The babble was presented continuously so that it was not time-locked to the stimulus, providing a constant backdrop of interference in the noise condition (e.g., Alain et al., 2012; Bidelman, 2016; Bidelman and Howell, 2016). Comparing behavioral performance between clear and degraded stimulus conditions allowed us to assess the impact of acoustic noise and differences between normal and hearing-impaired listeners in speech perception. Importantly, our task ensured that FFRs/ERPs were recorded online, during *active* speech perception. This helps circumvent issues in interpreting waveforms recorded across different attentional states or task demands (for discussion, see Bidelman, 2015a).

Stimulus presentation was controlled by a MATLAB (The Mathworks, Inc.; Natick, MA) routed to a TDT RP2 interface (Tucker-Davis Technologies; Alachua, FL) and delivered binaurally through insert earphones (ER-3; Etymotic Research; Elk Grove Village, IL). The speech stimuli were presented at an intensity of 75 dB_A_ SPL (noise at 65 dB_A_ SPL) using alternating polarity and FFRs/ERPs were derived by summing an equal number of condensation and rarefaction responses. This approach helps minimize stimulus artifact and cochlear microphonic from scalp recordings (which flip with polarity) and accentuates portions of the FFR related to signal envelope, i.e., fundamental frequency (F0) (Aiken and Picton, 2008; Skoe and Kraus, 2010b; Smalt et al., 2012).

### QuickSIN test

We measured listeners’ speech reception thresholds in noise using the QuickSIN test (Killion et al., 2004). Participants were presented lists of six sentences with five key words per sentence embedded in four-talker babble noise. Sentences were presented at 70 dB SPL using pre-recorded SNRs that decreased in 5 dB steps from 25 dB (very easy) to 0 dB (very difficult). Listeners scored one point for each key word correctly repeated. “SNR loss” (in dB) was determined as the SNR required to correctly identify 50% of the key words (Killion et al., 2004). SNR loss reflects the performance in noise compared to normal-hearing persons’ performance in noise. Consequently, larger scores reflect worse performance in SIN recognition. We averaged SNR loss from four list presentations per listener.

### Electrophysiological recordings and analysis

#### EEG acquisition and preprocessing

During the primary behavioral task, neuroelectric activity was recorded from 32 channels at standard 10-20 electrode locations on the scalp (Oostenveld and Praamstra, 2001). Recording EEGs during the active listening task allowed us to control for attention and assess the relative influence of brainstem and cortex during online speech perception. The montage included electrode coverage over frontocentral (Fz, Fp1/2, F3/4, F7/8, F9/10, C3/4), temporal (T7/8, TP7/9, TP8/10), parietal (Pz, P3/4, P7/8), and occipital-cerebellar (Oz, O1/2, CB1/2, Iz) sites. Electrodes placed along the zygomatic arch (FT9/10) and the outer canthi and superior/inferior orbit of the eye (IO1/2, LO1/2) monitored ocular activity and blink artifacts. Electrode impedances were maintained at ≤ 5 kΩ. EEGs were digitized at a sampling rate of 20 kHz using SynAmps RT amplifiers (Compumedics Neuroscan; Charlotte, NC). Data were re-referenced off-line to a common average reference for further analyses.

Subsequent pre-processing was performed in BESA^®^ Research v6.1 (BESA, GmbH). Ocular artifacts (saccades and blinks) were first corrected in the continuous EEG using a principal component analysis (PCA) (Picton et al., 2000). Cleaned EEGs were then epoched (-10-200 ms), baseline corrected to the pre-stimulus period, and subsequently averaged in the time domain to obtain compound evoked responses, containing both brainstem and cortical activity (Bidelman et al., 2013), for each stimulus condition per participant.

#### Source waveform derivations

Scalp potentials (sensor-level recordings) were transformed to source space using BESA. We seeded three dipoles located in (i) midbrain of the brainstem (BS) and (ii-iii) bilateral primary auditory cortex (PAC) (Bidelman, 2018). Dipole orientations for the PAC sources were set using the tangential component of BESA’s default auditory evoked potential (AEP) montage (Scherg et al., 2002). The tangential component was selected given that it dominantly explains the auditory cortical ERPs (Picton et al., 1999). Orientation of the BS source followed the oblique, fronto-centrally directed dipole of the FFR (Bidelman, 2015b). Focusing on BS and PAC source waveforms allowed us to reduce the dimensionality of the scalp data from 32 sensors to 3 source channels and allowed specific hypothesis testing regarding hearing-induced changes in brainstem-cortical connectivity. While simplistic, this model’s average goodness of fit (GoF) across groups and stimuli was 88.1±3.8%, meaning that residual variance (RV) between recorded and source-modeled data was low (RV= 11.9±3.9%).

To extract individuals’ source waveforms within each region of interest (ROI), we transformed their scalp recordings into source-level responses using a virtual source montage (Scherg et al., 2002). This digital re-montaging applies a spatial filter to all electrodes (defined by the foci of our three-dipole configuration). Relative weights were optimized in BESA to image activity within each brain ROI while suppressing overlapping activity stemming from other active brain regions (for details, see Scherg and Ebersole, 1994; Scherg et al., 2002). For each participant, the model was held fixed and was used as a spatial filter to derive their source waveforms (Alain et al., 2009; Zendel and Alain, 2014), reflecting the neuronal current (in units nAm) as seen *within* each anatomical ROI. Compound source waveforms were then bandpass filtered into high (100–1000 Hz) and low (1-30 Hz) frequency bands to isolate the periodic brainstem FFR vs. slower cortical ERP waves from each listeners’ compound evoked response (Musacchia et al., 2008; Bidelman et al., 2013; Bidelman, 2015a). Comparing FFR and ERP source waveforms allowed us to assess the relative contributions of brainstem and cortical activity to SIN comprehension in normal and hearing-impaired listeners. Results reported herein were collapsed across /ba/ and /pa/ tokens to reduce the dimensionality of the data. Infrequent /ta/ responses were not analyzed given the limited number of trials for this condition and to avoid mismatch negativities in our analyses.

#### FFR source waveforms

We measured the magnitude of the source FFR F0 to quantify the degree of neural phase-locking to the speech envelope rate, a neural correlate of “voice pitch” encoding (Bidelman and Krishnan, 2010; Parbery-Clark et al., 2013; Bidelman and Alain, 2015). F0 was the most prominent spectral component in FFR spectra (see Fig. 4) and is highly replicable both within and between listeners (Bidelman et al., 2018). F0 was taken as the peak amplitude in response spectra nearest the 150 Hz bin, the expected F0 based on our speech stimuli.

#### ERP source waveforms

Prominent components of the ERP source responses were quantified in latency and amplitude using BESA’s automated peak analysis for both left and right PAC waveforms in each participant. Appropriate latency windows were first determined by manual inspection of grand averaged traces. For each participant, the P1 wave was then defined as the point of maximum upward deflection from baseline between 40 and 70 ms; N1 as the negative-going deflection within 90 and 145 ms; P2 as the maximum positive deflection between 145 and 175 ms (Hall, 1992). These measures allowed us to evaluate the effects of noise and hearing loss on the magnitude and efficiency of cortical speech processing. Additionally, differentiation between hemispheres enabled us to investigate the relative contributions of each auditory cortex to SIN processing.

### Functional connectivity

We measured causal (directed) information flow between nodes of the brainstem-cortical network using phase transfer entropy (PTE) (Lobier et al., 2014). For data reduction purposes, responses were collapsed across left and right hemispheres and stimuli prior to connectivity analysis. PTE is a non-parametric, information theoretic measure of directed signal interaction. It is ideal for measuring functional connectivity between regions because it can detect nonlinear associations between signals and is robust against the volume conducted cross-talk in EEG (Vicente et al., 2011; Hillebrand et al., 2016). PTE was estimated using the time series of the instantaneous phases of pairwise signals (i.e., BS and PAC waveforms) (Lobier et al., 2014; Hillebrand et al., 2016). PTE was computed according to Eq. 1:

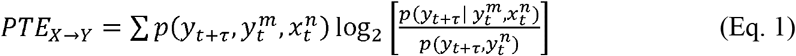

where *X* and *Y* are the ROI signals and the log(.) term is the conditional probabilities between signals at time *t+τ* for sample *m* and *n*. The probabilities were obtained by building histograms of occurrences of pairs of phase estimates in the epoch (Lobier et al., 2014). Following Hillebrand et al. (2016), the number of histogram bins was set to *e*^0.626+0.4ln*(Ns – τ –* 1)^ (Otnes and Enochson, 1972). The prediction delay τ was set at 100 ms, to include coverage of the entirety of the FFR signal and early cortical ERPs (see Fig. 2). Although this τ was based on *a priori* knowledge of the ERP time course, it should be noted that PTE yields similar results with comparable sensitivity across a wide range of analysis lags (Lobier et al., 2014).

**Figure 2:**
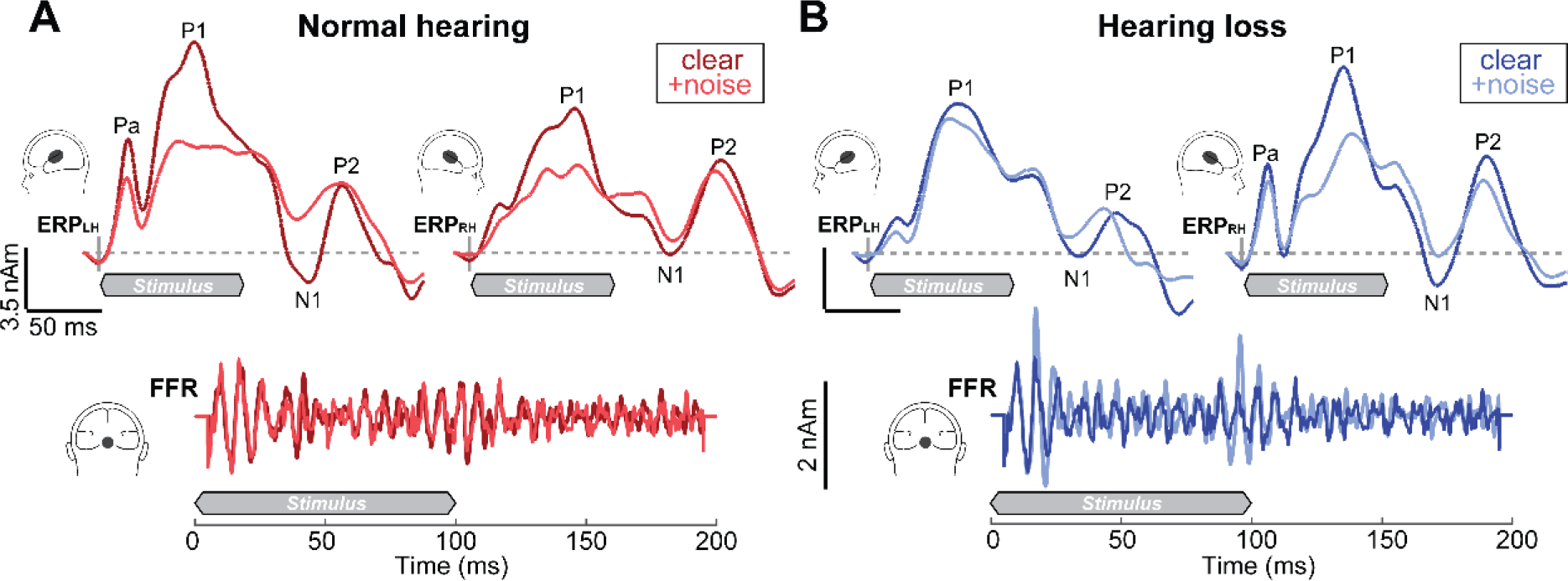
ERP (*top traces*) and FFR (*bottom*) source waveforms reflect the simultaneous encoding of speech within cortical and brainstem tiers of the auditory system. (A) NH listeners show a leftward asymmetry in PAC responses compared to HL listeners (B), who show stronger activation in right PAC. Noise weakens the cortical ERPs to speech across the board, particularly in the timeframe of P1 and N1, reflecting the initial registration of sound in PAC. In contrast to cortical responses, BS FFRs are remarkably similar between groups and noise conditions. Shaded regions demarcate the 100 ms speech stimulus. BS, brainstem; PAC, primary auditory cortex.

Intuitively, PTE is understood as the reduction in information (units bits) necessary to describe the present ROI_Y_ signal using both the past of ROI_X_ and ROI_Y_. PTE cannot be negative and has no upper bound. Higher values indicate stronger connectivity, whereas *PTE_X→Y_* =0 implies no directed signaling. In this sense, it is similar to the definition of Granger Causality (Barnett et al., 2009), which states that ROI_X_ has a causal influence on the target ROI_Y_ if knowing the past of both signals improves the prediction of the target’s future compared to knowing only its past. Yet, PTE has several important advantages over other connectivity metrics (Lobier et al., 2014): (i) PTE is more robust to realistic amounts of noise and linear mixing in the EEG that can produce false-positive connections; (ii) PTE relaxes assumptions about data normality and is therefore model-free; (iii) PTE is asymmetric so it can be computed bi-directionally between pairs of sources (X→Y vs. Y→X) to infer causal, directional flow of information between interacting brain regions. Computing PTE in both directions between BS and PAC allowed us to quantify the relative weighting of information flowing between subcortical and cortical ROIs in both feedforward (afferent; BS→PAC) and feedback (efferent; PAC→BS) directions.

### Statistical analysis and Experimental Design

Unless otherwise noted, two-way mixed model ANOVAs were conducted on all dependent variables (GLIMMIX, SAS^®^ 9.4, SAS Institute; Cary, NC). Group (2 levels; NH, HL) and stimulus SNR (2 levels; clear, noise) functioned as fixed effects; participants served as a random factor. With the exception of ERP amplitude measures (see below), initial diagnostics confirmed normality and homogeneity of variance assumptions for parametric statistics. Tukey–Kramer adjustments controlled Type I error inflation. An *a priori* significance level was set at α = 0.05 for all statistical analyses. Effect sizes are reported as Cohen’s-*d* (Wilson, 2018). Independent samples *t*-tests (un-pooled variance, two-tailed) were used to contrast demographic variables.

Correlational analyses (Pearson’s-*r*) and robust regression (bisquare weighting) were used to evaluate relationships between neural and behavioral measures. Robust fitting was achieved using the ‘fitlm’ function in MATLAB. We used an efficient, bootstrapping implementation of the Sobel statistic (Sobel, 1982; Preacher and Hayes, 2004) (*N*=1000 resamples) to test for mediation effects between demographic, neural connectivity, and behavioral measures.

## RESULTS

### Behavioral data

Behavioral accuracy and reaction time for target speech detection are shown for each group and noise condition in Figure 1. An ANOVA revealed a main effect of SNR on /ta/ detection accuracy, which was lower for noise-degraded compared to clear speech [*F*_1,30_=5.66, *p*=0.024, *d*=0.88; Fig. 1B]. However, groups differed neither in their accuracy [*F*_1,30_=0.01, *p*=0.94*; d*=0.04] nor speed [F_1,30_=0.47, *p*=0.49; *d*=0.26; Fig. 1C] of speech identification. On average, HL individuals achieved QuickSIN performance within ~1 dB of NH listeners, and scores did not differ between groups [*t*_2.35_=-1.43, *p*=0.16] (Fig. 1D). Nevertheless, HL listeners showed more inter-subject variability in SIN performance compared to NH listeners [Equal variance test (two-sample *F*-test): *F*_18,12_=8.81, *p*=0.0004]. Collectively, these results suggest that the hearing loss in our sample was not yet egregious enough to yield substantial deficits in speech perception.

### Electrophysiological data

Speech-evoked brainstem FFR and cortical ERP source waveforms are shown in Figure 2. Cortical activity appeared as a series of obligatory waves developing over ~200 ms after the initiation of speech that were modulated by noise and cerebral hemisphere. Noise-related changes in the ERPs were particularly prominent in the earlier P1 and N1 deflections reflecting the initial registration of sound in medial portions of PAC and secondary auditory cortex (Scherg and von Cramon, 1986; Liégeois-Chauvel et al., 1994; Picton et al., 1999).

These observations were confirmed via quantitative analysis of source ERP latency and amplitude (Fig. 3). ANOVA diagnostics indicated positive skew in ERP amplitude measures. Thus, we used a natural log transform in analyses of the cortical amplitude data. An ANOVA conducted on log-transformed ERP amplitudes revealed a main effect of SNR for both P1 and N1 with stronger responses for clear compared to noise-degraded speech [P1 amp: *F*_1, 94_=12.67, *p*<0.001, *d*=1.28; N1 amp: *F*_1, 94_=6.70, *p*=0.01, *d*=0.93; data not shown]. These results replicate the noise-related degradation in speech-evoked activity observed in previous studies (e.g., Alain et al., 2014; Bidelman and Howell, 2016). Unlike the early ERP waves, P2 amplitude varied between hemispheres [*F*_1,94_=9.38, *p*=0.003, *d*=1.10], with greater activation in right PAC. There was also a main effect of group with larger P2 responses in NH listeners [*F*_1,30_=4.74, *p*=0.038*, d*=0.78] (Fig. 3A and 3B). The P2 deflection is thought to reflect the signal’s identity, recognition of perceptual objects, and perceptual-phonetic categories of speech (Wood et al., 1971; Eulitz et al., 1995; Alain et al., 2007; Bidelman et al., 2013; Bidelman and Lee, 2015; Bidelman and Yellamsetty, 2017). The effects of age and noise on the P2 wave could indicate deficits in mapping acoustic details into a more abstract phonemic representation.

**Figure 3:**
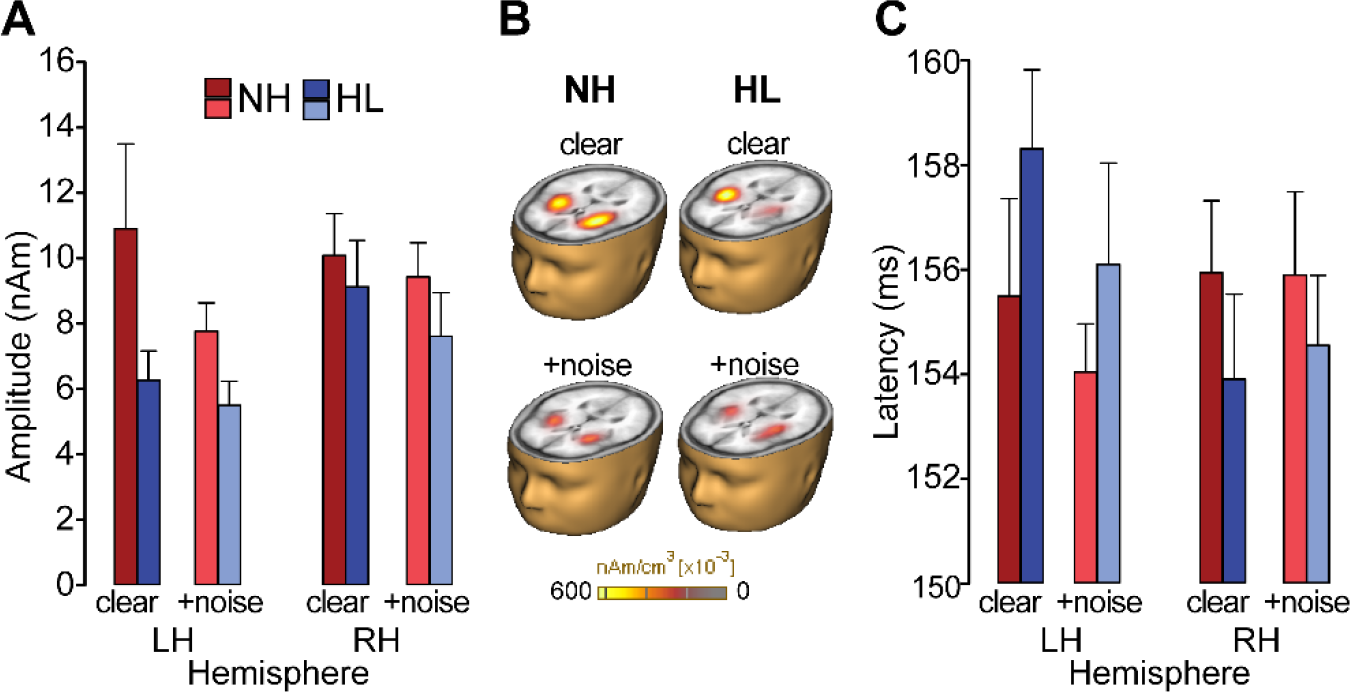
Cortical speech processing is modulated by noise interference, hearing status, and cerebral hemisphere. (A) P2 amplitudes are stronger in NH listeners regardless of SNR. (B) Brain volumes show distributed source activation maps using Cortical Low resolution electromagnetic tomography Analysis Recursively Applied (CLARA; BESA v6.1) (Iordanov et al., 2014). Functional data are overlaid on the BESA brain template (Richards et al., 2016). (C) P2 latency revealed a group x hemispheric interaction. In HL listeners, responses were ~3 ms earlier in right compared to left hemisphere (R<L) whereas no latency differences were observed in NH ears (L=R). errorbars = ± s.e.m.

For latency, no effects were observed at P1. However, hemispheric differences were noted for N1 latencies [*F*_1,94_=9.49, *p*=0.003*, d*=1.11], where responses were ~4 ms earlier in the right compared to left hemisphere across both groups. P2 latency also showed a group x hemisphere interaction [*F*_1,93_=5.27, *p*=0.02, *d*=0.82] (Fig. 3B). Post hoc analyses revealed a significant asymmetry for the HL group: P2 latencies were ~3 ms earlier in right relative to left PAC whereas no hemispheric asymmetry was observed in the NH listeners. These results indicate an abnormal hemispheric asymmetry beginning as early as N1 extending through P2 (~150 ms) in listeners with mild hearing impairment.

In contrast to slow cortical activity, brainstem FFRs showed phase-locked neural activity to the periodicities of speech (Fig. 2, bottom traces). Analysis of response spectra revealed strong energy at the voice fundamental frequency (F0) and weaker energy tagging the upper harmonics of speech (Fig. 4). Previous FFR studies have shown that older adults have limited coding of the high-frequency harmonics of speech (e.g., Anderson, Parbery-Clark, et al., 2013; Bidelman, Villafuerte, et al., 2014; Clinard and Cotter, 2015; Bidelman et al., 2017). The latter is particularly susceptible to noise (Bidelman and Krishnan, 2010; Bidelman, 2016) and hearing loss (Henry and Heinz, 2012) and reduced amplitudes may be attributable to age- and hearing-related changes in brainstem phase-locking (Parthasarathy et al., 2014). Weaker harmonic energy of the F0 may also be due to the relatively short duration of vowel periodicity (< 40 ms) of our stimuli. Group and noise-related effects in FFRs were less apparent than in the ERPs. An ANOVA conducted on FFR F0 amplitudes showed that FFR in older adults was little affected by hearing loss [main effect of group: *F*_1,30_=0.38, *p*=0.54, *d*=0.22] or background noise [main of effect of SNR: *F*_1,30_=0.41, *p*=0.53, *d*=0.23] (Fig. 4B). These results suggest that neither the severity of noise nor mild hearing impairment had an appreciable effect on the fidelity of brainstem F0 coding in our listeners. Yet, comparing across levels of the neuroaxis, age-related hearing loss had a differential effect on complex sound coding across levels, exerting a stronger effect at cortical vs. subcortical stages of the auditory system (cf. Bidelman, Villafuerte, et al., 2014).

**Figure 4:**
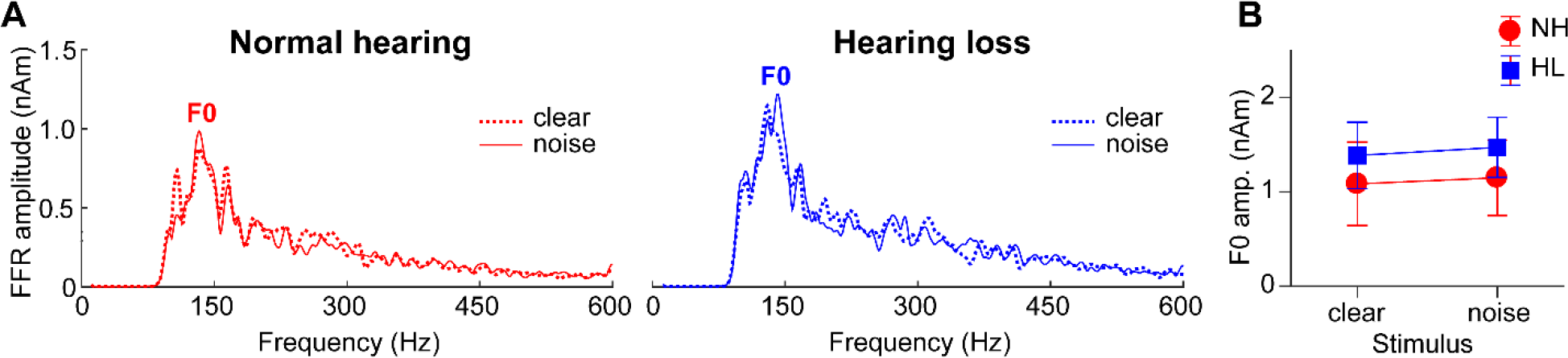
Brainstem speech processing as a function of noise and hearing loss. (A) Source FFR spectra for response to clear and degraded speech. Strong energy is observed at the voice fundamental frequency (F0) but much weaker energy tagging the upper harmonics of speech, consistent with age-related declines in high-frequency spectral coding. Group and noise-related effects in FFRs were less apparent than in the cortical ERPs (cf. Fig. 3). errorbars = ± s.e.m.

### Brainstem-cortical functional connectivity

Phase transfer entropy, quantifying the feedforward (afferent) and feedback (efferent) functional connectivity between BS and PAC, is shown in Figure 5. We found that afferent BS→PAC signaling was stronger in NH vs. HL listeners [*F*_1,30_=5.52, *p*=0.0256, *d*=0.84] (Fig. 5A) and negatively correlated with the degree of listeners’ hearing impairment based on their PTAs (Fig. 5B) [*r=-*0.59, *p*=0.0004]. Individuals with poorer hearing acuity showed reduced neural signaling directed from BS to PAC. More interestingly, we found afferent connectivity also predicted behavioral QuickSIN scores (Fig. 5C) [*r=-*0.65, *p*<0.0001], such that listeners with weaker BS→PAC transmission showed poorer SIN comprehension (i.e., higher QuickSIN scores)^1^.

**Figure 5:**
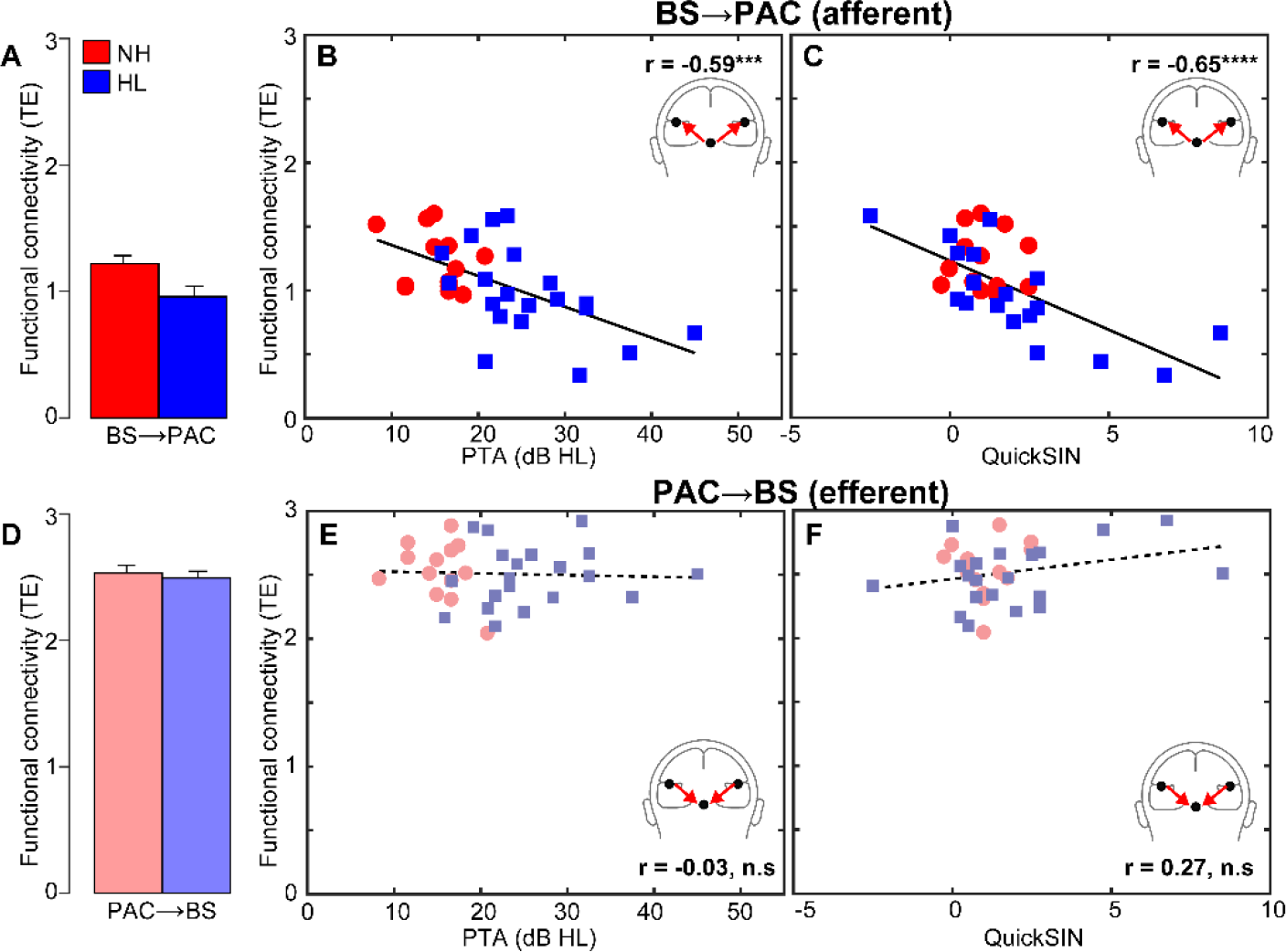
Functional connectivity between auditory brainstem and cortex varies with hearing loss and predicts SIN comprehension. (A) Transfer entropy reflecting directed (casual) *afferent* neural signaling from BS→PAC. Afferent connectivity is stronger in normal compared to hearing-impaired listeners. (B) Afferent connectivity is weaker in listeners with poorer hearing (i.e., worse PTA thresholds) and predicts behavioral SIN performance (C). Individuals with stronger BS→PAC connectivity show better (i.e., lower) scores on the QuickSIN. (D) *Efferent* neural signaling from PAC→BS does not vary between NH and HL listeners, suggesting similar top-down processing between groups. Similarly, efferent connectivity did not covary with hearing loss (E) nor did it predict SIN comprehension (F). Solid lines=significant correlations; dotted lines=*n.s.* relationships. errorbars = ± s.e.m., \*\*\**p*<0.001, \*\*\*\**p*<0.0001.

In contrast to afferent flow, efferent connectivity directed from PAC→BS, did not differentiate groups [*F*_1,30_=0.21, *p*=0.65, *d*=0.16] (Fig. 5D). Furthermore, while efferent connectivity was generally stronger than afferent connectivity [*t*_31_=2.52, *p*=0.0171], PAC→BS transmission was not correlated with hearing thresholds (Fig. 5E) [*r=-*0.03, *p*=0.86] nor behavioral QuickSIN scores (Fig. 5F) [*r=*0.27, *p*=0.14]. Collectively, connectivity results suggest that mild hearing loss alters the afferent-efferent balance of neural communication between auditory brainstem and cortical structures. However, in the aging auditory system, bottom-up (BS→PAC) transmission appears more sensitive to peripheral hearing loss (as measured by pure tone thresholds) and is more predictive of perceptual speech outcomes than top-down signaling (PAC→BS).

BS→PAC connectivity was correlated with both mild hearing loss and behavioral QuickSIN measures, which suggests that neural signaling could mediate SIN comprehension in older adults in addition to peripheral hearing loss. To test this possibility, we used Sobel mediation analysis (Sobel, 1982; Preacher and Hayes, 2004) to tease apart the contributions of hearing loss (PTA) and afferent connectivity (PTE) on listeners’ QuickSIN scores (among the entire sample). The Sobel test contrasts the strength of regression between a pairwise vs. a triplet (mediation) model (i.e., *X→Y* vs. *X*→*M*→*Y*). *M* is said to completely mediate the relation between the *X*→*Y* if (i) X first predicts Y on its own, (ii) X predicts *M*, and (iii) the functional relation between *X*→*Y* is rendered insignificant after controlling for the mediator *M* (Baron and Kenny, 1986; Preacher and Hayes, 2004).

PTA by itself was a strong predictor of QuickSIN scores (Fig. 6A) [*b*=0.13; *t*=3.23, *p*=0.0030]; reduced hearing acuity was associated with poorer SIN comprehension. However, when introducing BS→PAC afferent connectivity into the model, the direct relation between PTA and QuickSIN was no longer significant (Fig. 6B) [Sobel mediation effect: *z=*2.42, *p*=0.016]. PTA predicted the strength of afferent connectivity [*b*=-0.02; *t*=-4.01, *p*=0.0004] and in turn, connectivity predicted QuickSIN scores [*b*=-3.28; *t*=-3.12, *p*=0.0041], but the effect of hearing loss on SIN comprehension was indirectly mediated by BS→PAC connectivity strength^2^. In contrast to afferent signaling, efferent connectivity was not a mediator of SIN comprehension [Sobel *z=*-0.16, *p*=0.87]. However, this result might be anticipated given the lack of group differences in efferent PAC→BS connectivity. These results indicate that while hearing status is correlated with perception, the underlying afferent flow of neural activity from BS→PAC fully mediates older adults’ SIN listening skills.

**Figure 6:**
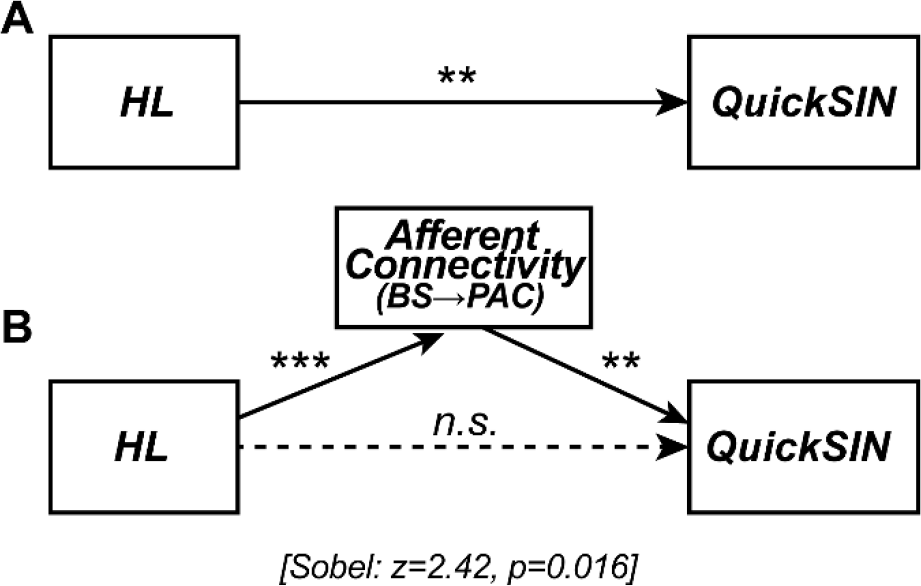
Afferent neural signaling from BS to PAC fully mediates the relation between hearing loss and SIN comprehension. Sobel mediation analysis (Sobel, 1982) between listeners’ hearing loss (PTA thresholds), neural connectivity (BS→PAC signaling), and SIN comprehension (QuickSIN scores). Edges show significant relations between pairwise variables identified via linear regression. (A) Hearing loss by itself strongly predicts QuickSIN scores such that reduced hearing is associated with poorer SIN comprehension. (B) Accounting for BS→PAC afferent connectivity renders this relation insignificant (Sobel test: z=2.42, p=0.016; Sobel, 1982), indicating the strength of neural communication between BS and PAC,

## DISCUSSION

By examining functional connectivity between brainstem and cortical sources of neuroelectric responses to speech, we demonstrate a critical dissociation in how hearing loss impacts neural sound representations and the transfer of information between functional levels of the auditory pathway. We show that afferent (BS→PAC), but not efferent (PAC→BS), neural transmission during active speech perception weakens with declining hearing and this connectivity predicts listeners’ SIN comprehension/identification. These findings reveal that while age-related hearing loss alters neural output *within* various tiers of the auditory system (PAC>BS) (i) bottom-up subcortical-cortical connectivity is more sensitive to diminished hearing than top-down (cortical-subcortical) connectivity, and (ii) afferent BS→PAC neural transmission accounts for reduced speech understanding in the elderly.

### Hearing loss differentially alters subcortical vs. cortical auditory processing

Comparisons between source-level FFRs and ERPs revealed that age-related hearing loss had a differential impact on brainstem vs. cortical speech processing. This finding is reminiscent of animal work demonstrating that online changes in inferior colliculus receptive fields are smaller and in the opposite direction of changes in auditory cortex for the same task (Slee and David, 2015). In our own EEG studies, we showed that hearing loss weakens brainstem encoding of speech (e.g., F0 pitch and formant cues) whereas both age and hearing loss exert negative effects at the cortical level (Bidelman, Villafuerte, et al., 2014). Here, we show that age-related hearing loss reduces amplitude and prolongs the latency of cortical speech activation, indicative of weaker and less efficient neural processing. In contrast, FFRs showed negligible group differences. The lack of significant difference related to age-related hearing loss in lower-level (BS) compared to higher-level (PAC) auditory sources suggests that declines in hearing acuity during the aging process exert a differential effect on neural encoding across functional stages of the auditory hierarchy. Our findings contrast those of prior FFR studies on aging (e.g., Anderson, Parbery-Clark, et al., 2013; Bidelman, Villafuerte, et al., 2014; Clinard and Cotter, 2015; Bidelman et al., 2017). The discrepancy may be due to the fact that our FFR analyses focused on source responses—a more “pure” measurement of midbrain activity—rather than scalp potentials (previous studies), which can blur the contributions of various subcortical and cortical FFR generators (Coffey et al., 2016; Bidelman, 2018). Moreover, we have found that changes in speech-FFRs only become apparent when hearing impairments exceed PTAs of 30-40 dB HL (Bidelman, Villafuerte, et al., 2014), which are greater than those observed in the present study. All the same, our results bolster the notion that brainstem and cortical mechanisms provide functionally distinct contributions to speech coding (Bidelman et al., 2013) and are differentially susceptible to the various insults of the aging process (Bidelman, Villafuerte, et al., 2014; Bidelman et al., 2017).

The lines between peripheral vs. central function and impaired sensory encoding vs. signal transmission issues are difficult to disentangle in humans (Humes, 1996; Marmel et al., 2013; Bidelman, Villafuerte, et al., 2014). Functional changes may result from an imbalance of excitation and inhibition in brainstem (Parthasarathy and Bartlett, 2012), cortex (Chao and Knight, 1997; Caspary et al., 2008), or both structural levels (Bidelman, Villafuerte, et al., 2014). Conversely, neurodegeneration at peripheral sites may partially explain our findings (Makary et al., 2011). Under this interpretation, observed changes in midbrain FFR and cortical PAC activity might reflect maladaptive plasticity in response to deficits earlier (lower) in the pathway. However, we would expect that degeneration due to age alone would produce similar effects between groups since both cohorts were elderly listeners. Instead, it is likely that listeners’ hearing loss (whether central or peripheral in origin) is what produces the cascade of functional changes that alter the neural encoding of speech at multiple stages of the auditory system. In this sense, our data corroborate *in vivo* evidence in animals that central (cortical) gain helps restore diminished sensory input (cf. brainstem) following cochlear damage (Chambers et al., 2016). Interestingly, such peripheral-induced neural rebound is stronger at cortical compared to brainstem levels (Chambers et al., 2016), consistent with the more extensive changes we find in human PAC relative to BS responses. Our data are also consistent with the notion that complex sound representations at peripheral sites (i.e., brainstem) are more affected by noise than their corresponding cortical representations (Rabinowitz et al., 2013; Bidelman et al., in press).

Our ERP data further imply that hearing loss might reorganize functional asymmetries at the cortical level (Du et al., 2016; Pichora-Fuller et al., 2017). Source waveforms from left and right PAC revealed that the normal hearing listeners showed bilateral symmetric cortical activity (Figs. 2-3). This pattern was muted in listeners with mild hearing impairment, who showed faster response in right hemisphere. These differences imply that the hemispheric laterality of speech undergoes a functional reorganization following sensory loss where processing might be partially reallocated to right hemisphere in a compensatory manner. Similar shifts in the cortical activity have been observed in sudden onset, idiopathic hearing loss (He et al., 2015), implying that our results might reflect central reorganization following longer-term sensory declines. Previous studies have also shown that hemispheric asymmetry is correlated with SIN perception (Javad et al., 2014; Bidelman and Howell, 2016; Thompson et al., 2016). Conceivably, the reduction in left hemisphere speech processing we find in hearing-impaired listeners, along with reduced BS→PAC connectivity, might reflect a form of aberrant cortical function that could exacerbates SIN comprehension behaviorally.

Our cortical ERP data contrast recent reports on senescent changes in the cortical encoding of speech. Previous studies have shown larger ERP amplitudes to speech and non-speech stimuli among older relative to younger listeners (Herrmann et al., 2013; Bidelman, Villafuerte, et al., 2014; Presacco et al., 2016), possibly resulting from the peripheral auditory filter widening (Herrmann et al., 2013) and/or decreased top-down (frontal) gating of sensory information (Chao and Knight, 1997; Peelle et al., 2011; Bidelman, Villafuerte, et al., 2014). In contrast, studies reporting *larger* ERP amplitudes in older, hearing-impaired adults focus nearly entirely on scalp (i.e., electrode-level) responses, which mixes temporal and frontal source contributions that are involved in SIN processing in younger (Du et al., 2014; Bidelman and Dexter, 2015; Bidelman and Howell, 2016; Bidelman et al., in press) and especially older adults (Du et al., 2016). A parsimonious explanation of our ERP data then, is that weaker auditory cortical responses reflect reduced sensory encoding (within PAC) secondary to the diminished stimulus input from hearing loss.

### The critical role of brainstem-cortical connectivity for degraded speech perception

Our results extend previous brainstem and cortical studies by demonstrating age-related changes in the neural representations within certain auditory areas but also how information is communicated *between* functional levels. Notably, we found that robust feedforward neural transmission between brainstem and cortex is necessary for successful SIN comprehension in older adults, particularly those with mild hearing loss. To our knowledge, this is the first direct demonstration of auditory brainstem-cortical connectivity in humans and how this functional reciprocity relates to complex listening skills.

Despite ample evidence for online subcortical modulation in animals (Suga et al., 2000; Bajo et al., 2010; Slee and David, 2015; Vollmer et al., 2017), demonstrations of corticofugal effects in human brainstem responses have been widely inconsistent and loosely inferred through manipulations of task-related attention (Picton et al., 1971; Woods and Hillyard, 1978; Rinne et al., 2007; Skoe and Kraus, 2010a; Varghese et al., 2015; Forte et al., 2017). Theoretically, efferent modulation of brainstem should occur only for behaviorally relevant stimuli in states of goal-directed attention (Suga et al., 2002; Slee and David, 2015; Vollmer et al., 2017), and should be stronger in more taxing listening conditions (e.g., difficult SIN tasks; Krishnan and Gandour, 2009). In this regard, our assay of central connectivity during online SIN identification should have represented optimal conditions to detect possible afferent-efferent BS-PAC communication most relevant to behavior.

Our findings revealed that corticofugal (PAC→BS) efferent signaling was stronger than afferent connectivity overall, implying considerable top-down processing in older adults. These results converge with theoretical frameworks of aging that posit higher-level brain regions are recruited to aid speech perception in older adults (Reuter-Lorenz and Cappell, 2008; Wong et al., 2009). Behaviorally, older adults tend to expend more listening effort during SIN recognition than younger individuals (Gosselin and Gagne, 2011). Consequently, one interpretation of our data is that the elevated, invariant PAC→BS efferent connectivity we observe across the board reflects an increase in older adults’ listening effort or deployment of attentional resources. Such corticofugal engagement might enhance impoverished BS processing that normally declines with age (Anderson, White-Schwoch, et al., 2013b; Marmel et al., 2013; Bidelman, Villafuerte, et al., 2014; Clinard and Cotter, 2015; Bidelman et al., 2017), effectively normalizing its output and group differences in FFR responses (e.g., Fig. 4). However, we note that efferent connectivity was not associated with hearing loss or SIN performance, despite our use of an active listening task. Without concomitant data from younger adults (and passive tasks) it remains unclear how (if) the magnitude of corticofugal connectivity might change across the lifespan or with more egregious hearing impairments. Additionally, mild cognitive impairment is known to alter brainstem and cortical speech processing (Bidelman et al., 2017). As we did not measure cognitive function, it is possible that at least some of group differences we observe in BS→PAC connectivity reflect undetected cognitive decline, since auditory processing often covaries with cognitive function (Humes et al., 2013).

In stark contrast, afferent directed communication (BS→PAC) differentiated normal- and hearing-impaired listeners and was more sensitive to hearing loss than corticofugal signaling. More critically, afferent transmission was a strong predictor of listeners’ reduced speech understanding at the behavioral level and fully mediated speech-in-noise (QuickSIN) performance, above and beyond hearing loss, *per se*. Said differently, we found that afferent connectivity was necessary to explain the link between hearing loss (i.e., a marker of peripheral cochlear integrity) and SIN perception (behavior). Simplicity of our task notwithstanding, these neurophysiological changes in cross-regional communication seem to precede behavioral SIN difficulties since groups showed similar levels of performance in SIN detection despite neurological variations. This agrees with notions that sensory coding deficits in brainstem-cortical circuitry mark the early decline of hearing and other cognitive abilities resulting from biological aging or neurotrauma (Bidelman et al., 2017; Kraus et al., 2017).

Our data align with previous neuroimaging studies suggesting that age-related hearing loss is associated with reduced gray matter volume in auditory temporal regions (Eckert et al., 2012; Lin et al., 2014), PAC volume (Husain et al., 2011; Peelle et al., 2011; Eckert et al., 2012), and compromised integrity of auditory white matter tracts (Chang et al., 2004; Lin et al., 2008). Accelerated neural atrophy from hearing impairment is larger in right compared to left temporal lobe (Peelle et al., 2011; Lin et al., 2014). Such structural changes might account for the functional declines and redistribution of cortical speech processing among our hearing-impaired cohort. Diffusion tensor imaging also reveals weaker fractional anisotropy (implying reduced white matter) in the vicinity of inferior colliculus in listeners with sensorineural hearing (Lin et al., 2008). These structural declines in brainstem could provide an anatomical basis for the reduced functional connectivity (BS→PAC) among our hearing-impaired cohort.

Collectively, our findings provide a novel link between (afferent) subcortical-cortical *functional* connectivity and individual differences in auditory behavioral measures related to cocktail party listening (SIN comprehension). We speculate that similar individual differences in BS↔PAC connectivity strength might account more broadly for the pervasive and parallel neuroplastic changes in brainstem and cortical activity observed among highly experienced listeners, certain neuropathologies, and successful auditory learners (Musacchia et al., 2008; Chandrasekaran et al., 2012; Bidelman, Weiss, et al., 2014; Bidelman and Alain, 2015; Bidelman et al., 2017; Kraus et al., 2017; Reetzke et al., 2018). Our findings underscore the importance of brain connectivity in understanding the biological basis of age-related hearing deficits in real-world acoustic environments and pave the way for new avenues of inquiry into the biological basis of auditory skills.

## Acknowledgements

This work was supported by grants from the Canadian Institutes of Health Research (MOP 106619) and the Natural Sciences and Engineering Research Council of Canada (NSERC, 194536) awarded to C.A, and The Hearing Research Foundation awarded to S.R.A, and NIH/NIDCD R01DC016267 awarded to G.M.B.

## Author contributions

C.A., D.S., and S.A. designed the experiment; D.S. and S.A. collected the data; G.M.B and C.N.P. analyzed the data; all authors contributed to the interpretation of the results and the writing of the manuscript.

## Footnotes

1 An identical pattern of results was observed when considering correlations between listeners’ average audiometric thresholds (from 250-8000 Hz) which defined the NH and HL group membership (see Methods). BS →PAC afferent (but not efferent) connectivity was negatively correlated with average hearing thresholds (*r=-*0.63, *p*=0.0001; data not shown).

2 Although the causality would be questionable, we also could treat PTA as a mediator between afferent connectivity and QuickSIN scores (i.e., PTA *BS/PAC QuickSIN*). Importantly, this arrangement was not significant [Sobel *z=*-13.04, *p*=0.29]. This (i) indicates hearing loss (PTA) does not mediate the relation between afferent BS PAC connectivity and SIN and (ii) strengthens the causality of the relation between neural afferent signaling and QuickSIN performance reported in the text.

